# Preliminary Insights into the Effects of Spinal Manipulation Therapy of Different Force Magnitudes on Blood Biomarkers of Oxidative Stress and Pro-Resolution of Inflammation Mediators

**DOI:** 10.1101/2023.12.28.573549

**Authors:** Felipe C. K. Duarte, Martha Funabashi, David Starmer, Wania A. Partata

## Abstract

**Introduction:** Evidence has reported that spinal manipulation therapy (SMT) leads to spine segmental hypoalgesia through neurophysiological and peripheral mechanisms related to regulating inflammatory biomarker function. However, these studies also showed substantial inter-individual variability in the biomarker responses. Such variability may be due to the incomplete understanding of the fundamental effects of force-based manipulations (e.g., patient-specific force-time characteristics) on a person’s physiology in health and disease. This study investigated the short-term effects of distinct SMT force-time characteristics on blood oxidative stress and pro-resolution of inflammation biomarkers.

**Methods:** Nineteen healthy adults were clustered into three groups: control (preload only), target total peak force of 400N, and 800N. A validated force-sensing table technology (FSTT^®^) determined the SMT force-time characteristics. Blood samples were collected at pre-intervention, immediately after SMT, and 20 minutes post-intervention. Parameters of the oxidant system (total oxidant status, lipid peroxidation and lipid hydroperoxide), the antioxidant system (total antioxidant capacity and bilirubin), and lipid-derived resolvin D1 were evaluated in plasma and erythrocytes through enzyme-linked immunosorbent assay and colorimetric assays.

**Results:** Overall, 400N seemed to decrease blood oxidants, and 800N generally increased blood oxidants, decreased antioxidants and resolvin D1 mediator

**Conclusion:** Our findings indicate that different single SMT force-time characteristics presented contrasting effects on the systemic redox signalling biomarkers and pro-resolution of inflammation mediators in healthy participants, providing baseline information and direction for future studies in a clinical population.

## Introduction

Chronic musculoskeletal (cMSK) disorders, such as chronic spinal pain, are progressive, long-lasting, highly disabling disorders, representing a significant economic burden to the population and healthcare system worldwide.^1,2^ The World Health Organization (WHO) has recently released a management guideline for adults with chronic primary low back pain (CPLBP) recommending spinal manipulation therapy (SMT) as part of non-surgical interventions to help people experiencing CPLBP.^3^ Despite the recommendation, there is an incomplete understanding of the mechanism of SMT on health and chronic pain person physiology.

Spinal manipulation therapy is a force-based therapy modality that belongs to a class of treatments labelled as mechanotherapies.^4,5^ A recent review highlighted that SMT leads to spine segmental hypoalgesia through neurobiological mechanisms regulating inflammatory function.^6^ For instance, pre-clinical and clinical evidence demonstrated changes in systemic markers of inflammation and oxidative stress following SMT in people with and without neck and low back pain.^7–9^ Oxidative stress is recognized as free radical dyshomeostasis, meaning that elevated formation of free radicals damages proteins, lipids and DNA, challenging the cellular and, eventually, the system homeostasis.^10^ In contrast, at low physiological levels, they act as redox signalling molecules in essential cellular signalling pathways.^10^ Importantly, oxidative stress biomarkers have shown positive stratification and predictor value to several clinical conditions such as risk of cardiovascular disease, short-term acute ischemic stroke death, work stress and overwork in health care practitioners, coronary artery disease, and cognitive decline.^11–15^ In cMSK, oxidative stress has emerged as a plausible biomarker owing to its intrinsic molecular ability to affect the structure and function of the neuroimmune system with clinical implications on nociceptive amplification and diminished physical and cognitive function.^16,17^ Thus, targeting the production and detoxification of oxidants as a rehabilitative strategy may greatly benefit health and disease.

While it is known that SMT has modulatory effects on systemic markers of inflammation and oxidative stress biomarkers, the changes following SMT showed substantial inter-individual variability.^7,18,19^ This variability may be due to the lack of understanding of the fundamental effects of the mechanical loading of the SMT (e.g., patient-specific force-time characteristics) on a person’s neurophysiology. Recent preliminary evidence by our team exploring different SMT force-time characteristics and blood inflammatory biomarkers in healthy individuals observed a short-term increase of plasma pro-inflammatory IFN-g, IL-5, and IL-6 cytokines when thoracic SMT of total peak force around 800N was delivered.^4^ An opposite behaviour on these biomarkers was observed in SMT of lower peak forces.^4^ Although these findings should be taken with a grain of salt, especially regarding clinical relevance, they open an opportunity to continue this research line investigating further biological components of the neuroimmune function that are responsive to variations of SMT’s mechanical loading.

It is well-known that cytokines have an essential role in the inflammatory process; however, resolvins, a lipid-derived pro-resolution mediator, have emerged as a class of active mediators in counteracting inflammation.^20^ Resolvins regulate inflammatory and neuroinflammation processes by binding to their receptors on immune and neuronal cells, which eventually implicates dampening inflammation, nociception, sensitization, and pain perception.^20,21^ Thus, both resolvins and oxidative stress biomarkers hold the potential to be further studied, given their participation in cellular signalling, inflammation and peripheral and central sensitization.^16,17,21^ However, no previous research studied the relationship between mechanical loadings of SMT and pro-resolution mediator and biomarkers of oxidative stress. Therefore, this study aimed to explore the short-term effect of SMT with different force-time characteristics on systemic oxidative stress biomarkers and lipid-derived pro-resolution mediators in healthy individuals. We hypothesized that higher changes on these biomarkers would be seen in the short term when SMT of higher force magnitudes were applied, contrasting with lower SMT force magnitude. The findings from this exploratory study in young, healthy individuals will help inform future studies in the neurobiology and neuroimmunology related to mechanotherapy that can later be investigated in patients with chronic spine pain disorders such as CPLBP.

## Methods

### Design Overview

This parallel repeated-measures study was designed to capture potential treatment and time-dependent alterations of blood biomarkers of oxidative stress and pro-resolution mediators before and after a single thoracic SMT. This study was approved by the Canadian Memorial Chiropractic College (CMCC) Research Ethics Board (REB# 192034). All participants provided written informed consent before participating in this study.

### Study Participants

Adults of both sexes aged between 18 and 45 from CMCC were invited to participate through e-mail, posters, and word of mouth between February and March 2020 before COVID-19.

### Inclusion and Exclusion criteria

Potential participants were screened by a single examiner (FD) using a standardized screening form. Participants were included if they were asymptomatic (i.e., reporting no pain in any region of the body in the previous 30 days), had not received SMT in the last seven days, and did not present any acute health condition (e.g., acute musculoskeletal injury, cold, flu) in the week before the data collection day. Participants were excluded if they presented with central nervous system diseases (e.g., depression), inflammatory conditions (e.g., rheumatoid arthritis), hypertension, or chronic metabolic conditions (e.g., diabetes) that require regular intake of medication. Participants were excluded if they were pregnant or presented with any contraindication to thoracic SMT, such as a history of spinal surgery, thoracic spine fracture, spinal cord injury, osteoporosis, spinal infection, or neoplasm.

### Sample Size

The sample size was estimated based on a previous pre-post-experimental study. In contrast, SMT was applied to asymptomatic individuals, resulting in a reported effect size of 30% (average effect size) on plasma biomarkers^22^. By using the General Power Analysis Program (G*Power) (University of Trier, Germany), it was estimated that for a repeated-measures study with three groups and three measurement points, 9 participants per group (3 groups) would be required to achieve an effect size of 0.30, with an alpha error of 5%, and power of 80%. Accounting for a 20% drop-out rate, we adjusted the target sample size to 11 participants per group (33 total).

### Randomization and Allocation

Using a random number generator (Microsoft Excel), participants were randomly allocated by an investigator not involved in data collection (MF) to one of the three groups: control (preload force only), 400N (400 N total peak force), and 800N (800 N total peak force). Group allocation was done by concealing opaque envelopes containing the intervention groups’ and study participants’ identification numbers. The envelopes were handed to the chiropractor (DS) performing the SMT on the testing session day using consecutively opened envelopes. The participants were blinded to their group allocation and, consequently, the experimental group. The examiner assessing the outcome measures was blinded to the participant’s group allocation.

### Force time characteristics: Data acquisition

Analog data from the embedded force plate of the force-sensing table (FSTT^®^, Toronto, ON, Canada) were digitally sampled at 2000 Hz and converted to units of force (Newtons) using the manufacturer-specified calibration matrices. Preload force is defined as the force applied during the initial setup, the total peak force is designated as the maximum force value of the thrust, including the preload force, and time to peak (milliseconds) is defined as the time elapsed between preload force, Total peak force was extracted along all three axes of the force plate’s frame of reference using customized software (MATLAB, The MathWorks Inc., Natick, Massachusetts, USA). Given the posterior to anterior (P-A) nature of the forces applied during the thoracic SMT used in this study, variables from the P-A forces (forces vertical to the table - Fz) were extracted for SMT force-time characterization.

### Intervention

A P-A thoracic SMT was applied at the T6-T9 spine region by a chiropractor (DS) with experience with the FSTT^®^ to modulate the SMT force-time characteristics on pre-determined force targets. Two experimental groups received thoracic SMT with a total peak force magnitude of 400N (±150N) (group 400N) and with a total peak force magnitude of 800N (±150N) (group 800N). We chose 400N peak force since it represents the average total peak force applied during P-A thoracic SMT according to the literature.^23^ The 800N peak force was selected because it represents a two-fold peak force from the reported average, and it is a common pre-determined peak force target during educational training^23^. To ensure SMT was delivered with the pre-determined forces (400N and 800N), real-time visual feedback of SMT force-time graphs was provided to the chiropractor. A third group, receiving P-A thoracic preload force (without thrust), was designed as a control group.

### Blood sampling and treatment for analysis

Upon arrival at the laboratory, participants sat comfortably for 5 to 10 minutes while demographic characteristics (age, sex, height, weight) were recorded. At this point, participants were asked to confirm whether they still met the inclusion criteria, including no pain in any region of the body in the previous 30 days, no acute health issue in the last seven days and had not received SMT in the previous seven days. In addition, participants were asked to complete a set of visual analog scales (VAS) rating their pre-intervention discomfort levels at the thoracic shoulder or neck area region from 0 to 10, corresponding to no discomfort and maximum discomfort, respectively. Participants also rated their discomfort level (VAS) a second time immediately after intervention and for the third time 20 minutes after intervention.

Since we aimed to collect blood in three short-term time points (pre-, immediately after and 20 minutes after intervention), a peripheral intravenous line was established to avoid multiple venipunctures. Blood samples were drawn in commercially available 10mL tubes (vacutainers) containing EDTA anti-coagulant, gently shaken (2x) to mix blood with EDTA, and kept at room temperature until participants’ sample collections were completed. The initial 2mLs were obtained in a tube and discarded^24^. Blood was drawn with subjects seated at baseline, immediately after intervention, and 20 minutes after intervention. All samples from the study participants were completed in the morning (from 8 am to 12 pm) to avoid circadian rhythm influences on the biochemical parameters assessed^25^. After completing the three blood draws from each participant, blood samples were centrifuged at 3000 RPM for 15 minutes at four °C in a refrigerated centrifuge (Beckman Coulter Allegra X-22 Series). Plasma samples were aliquoted and stored at -80°C. Red blood cells (RBCs) were collected and prepared as described previously ^26^ for analyses of erythrocyte hydroperoxide content (a biomarker of oxidative stress)^24^. For all biochemical studies, a microplate reader capable of measuring absorbance from 200 to 999nm (Epoch absorbance reader, BioTek-VT-USA) was used; the assays were carried out in duplicate within ten months after collection.

### Oxidative stress biomarkers: Pro-oxidant parameters

### Total oxidant status (plasma)

Total oxidative status (TOS) in blood samples can be used to estimate the overall individual pro-oxidative state. TOS was determined using the colorimetric method described by Erel^27^. The TOS concentration was calculated from the calibration curve of hydrogen peroxide, a potent oxidant. Results regarding micromolar hydrogen peroxide equivalent per litre (μmol H2O2 Equiv/L) were expressed.

### Lipid peroxidation (LPO) (plasma)

Lipid peroxidation is a reactive process of oxidative stress-induced cell membrane damage by free radicals and reactive oxygen species (ROS). LPO is commonly determined by the quantification of its product, malondialdehyde (MDA). This study determined MDA content by the surrogate quantification of thiobarbituric acid reagents (TBARS)^28^. For the quantitative determination of TBARS, the R&D commercially available assay kit (TBARS assay, #KGE013) was used according to the manufacturer’s instructions. Results were expressed in micromolar of TBARS.

### Lipid Hydroperoxide content (LOOH) (erythrocytes)

Lipid hydroperoxide is a non-radical intermediate product generated by lipid peroxidation. The content of hydroperoxides in erythrocytes was determined using the method reported by Ochoa et al. (2003)^29^. Results were expressed in terms of micromolar per milligram of protein (μmol/mg protein).

### Oxidative stress biomarkers: Antioxidant parameters

### Total antioxidant capacity (TAC) (plasma)

TAC works as a surrogate measure of the function of a pool of enzymatic and nonenzymatic antioxidants in the plasma sample that counteract pro-oxidant products. We used the total antioxidant capacity (TAC) assay kit (Cell Meter™ Colorimetric Antioxidant Activity Assay Kit, # 15900) according to the manufacturer’s protocol (AAT Bioquest). The antioxidant capacity of the sample was compared with that of Trolox, a potent antioxidant, and it was calculated as Trolox millimolar (mM) equivalent per litre.

### Total Bilirubin (plasma)

According to the manufacturer’s protocol, plasma total bilirubin concentration was measured using a bilirubin assay kit (Sigma-Aldrich, #MAK126). Results are expressed in milligrams per deciliter (mg/dL).

### Pro-resolution mediator (Resolvin D1) (plasma)

Resolvin D1 is a specialized pro-resolving lipid mediator derived from omega-3 polyunsaturated fatty acids that play a crucial role in the resolution of inflammation and tissue repair. Resolvin D1 concentration was measured using a competitive ELISA kit per the manufacturer’s protocol (Cayman Chemist, MI-USA, #500380). Results are expressed in picograms per millilitre (pg/mL)

### Protein measurement

Protein was measured using Bradford’s method according to manufacturer orientation (Millipore-Sigma, #1103060500, ON-Canada).

## Outcomes

The primary outcome measures were blood parameters of oxidants, antioxidants, and pro-resolvin mediator (resolving D1) sampled at pre-intervention, immediately after the intervention, and 20 minutes after the intervention. Secondary outcomes included the level of discomfort (VAS) during, immediately and after 20 minutes of the intervention in the thoracic area, shoulder or neck area, and cavitation. The number of individuals presenting with discomfort relative to baseline in the VAS and the clinician’s perception of an audible popping sound (cavitation) after intervention was recorded and reported.

## Statistical Analysis

Participant characteristics (sex, age, weight, height, and body mass index calculation) were reported descriptively. Similarly, force-time characteristic outputs, presence of cavitation and number of participants with discomfort after intervention compared to baseline. Blood parameters of oxidants, antioxidants and resolvin D1 were analyzed using a mixed-effect model (REML) for small sample sizes to compare the participants’ biomarkers over time (baseline, immediately after, and after 20 minutes intervention) between the three groups (control, target 400, and 800N), followed by Tukey’s multiple comparison test.^30^ Pearson correlation coefficient was conducted to assess the linear relationship between the quantitative variables studied (i.e., force magnitude, oxidative stress biomarkers and pro-resolution mediator). Low (positive or negative) correlation corresponded to .30 to .50; moderate (positive or negative) corresponded to .50 to .70; high (positive or negative) correlation corresponded to .70 to 1.^31^

Data are presented as mean ± standard deviation (SD). Statistical analyses were performed in Prism (V. 9.0). Differences were considered statistically significant when the p-value was ≤ 0.05.

## Results

Due to the COVID-19 global pandemic, participant recruitment was limited to before the lockdown period between February and March 2020. Thirty-seven participants were identified and assessed for eligibility (Figure 1). Twenty-one participants met the inclusion criteria. Out of the 21, two were excluded: one due to fainting after the first phlebotomy and the other because the pre-established limit of peak force variability (±150N) was exceeded in the target 400 N. Therefore, data from 19 participants were used for the study analysis (Table 1).

**Figure 1:**
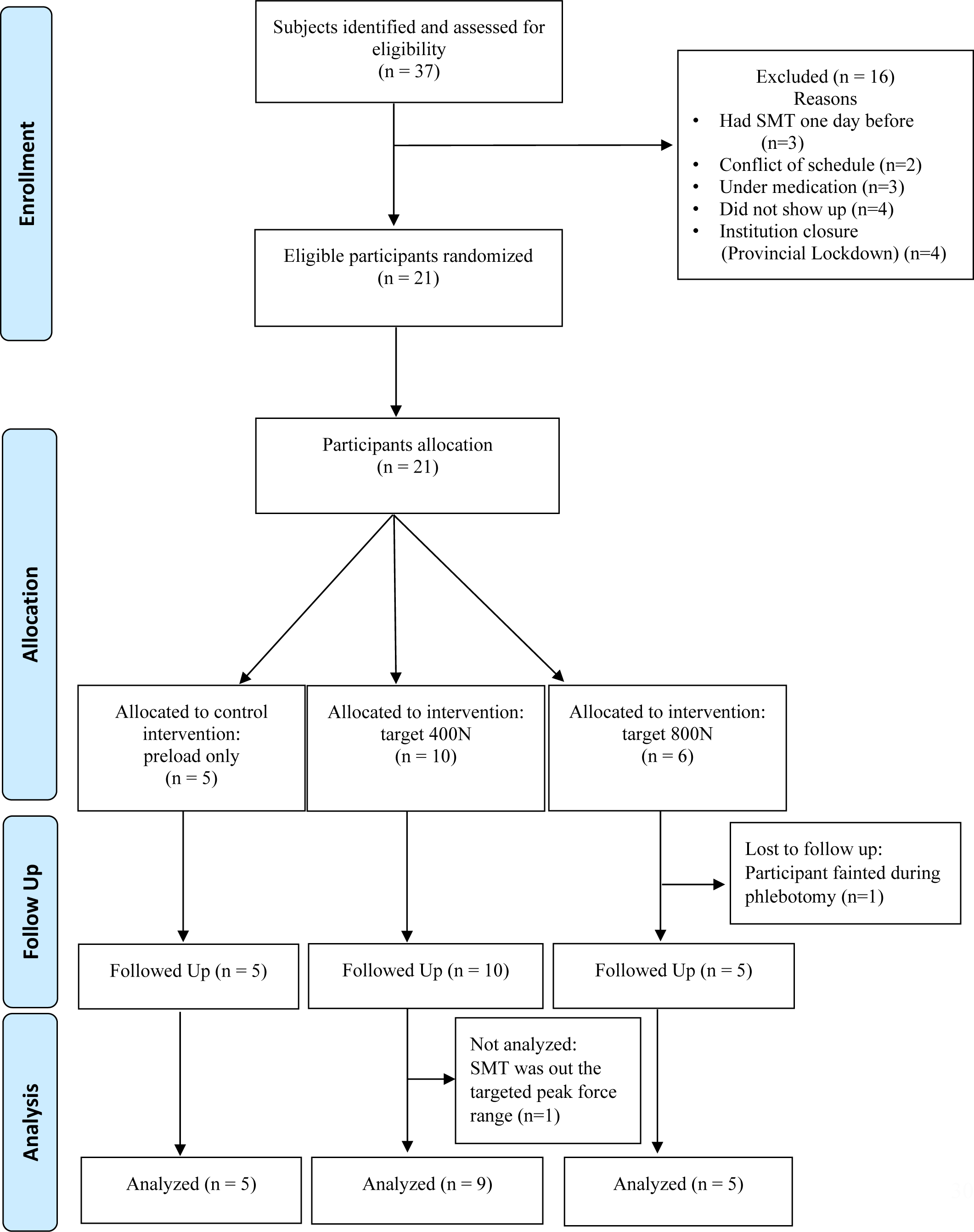
Study flow chart shows the number of participants for each intervention group included in the data analysis.

**Table 1:** Mean and standard deviation of demographic characteristics of the study participants.

| **Characteristics** | **Group Control**<br>**(n=5)** | **Group 400N**<br>**(n=9)** | **Group 800N**<br>**(n=5)** |
| --- | --- | --- | --- |
| Males: Females | 3:2 | 3:6 | 3:2 |
| Age (years) | 25.20 ± 0.83 | 25.30 ± 1.32 | 26.4 ± 1.35 |
| Weight (kg) | 66.80 ± 17.46 | 77.17 ± 16.48 | 67.00 ± 3.46 |
| Height (cm) | 172.60 ± 11.55 | 177.40 ± 10.37 | 171.30 ± 3.96 |
| Body mass index (kg/m<sup>2</sup>) | 22.07 ± 3.21 | 24.25 ± 2.70 | 22.86 ± 1.57 |

Kg: kilograms; cm: centimetres; m<sup>2</sup>: square meters.

### Preload, total peak force and time to peak

Preload values were comparable between groups ranging from 151N to 220N with the mean(±SD) of 202.4 (16.38), 183.4 (14.24) and 197.8 (9.86) in the control, target 400 and 800N, respectively (Table 2). Total peak force and time to peak were recorded only on the target 400N and 800N groups. The total peak force was highly consistent at the intended targets of 400 N or 800 N, with small variability between individuals within each group (Table 2). As expected, the time to peak was slightly faster in the target 400 N group, approximately 100ms, compared to 134ms in the target 800 N group (Table 2).

**Table 2:**
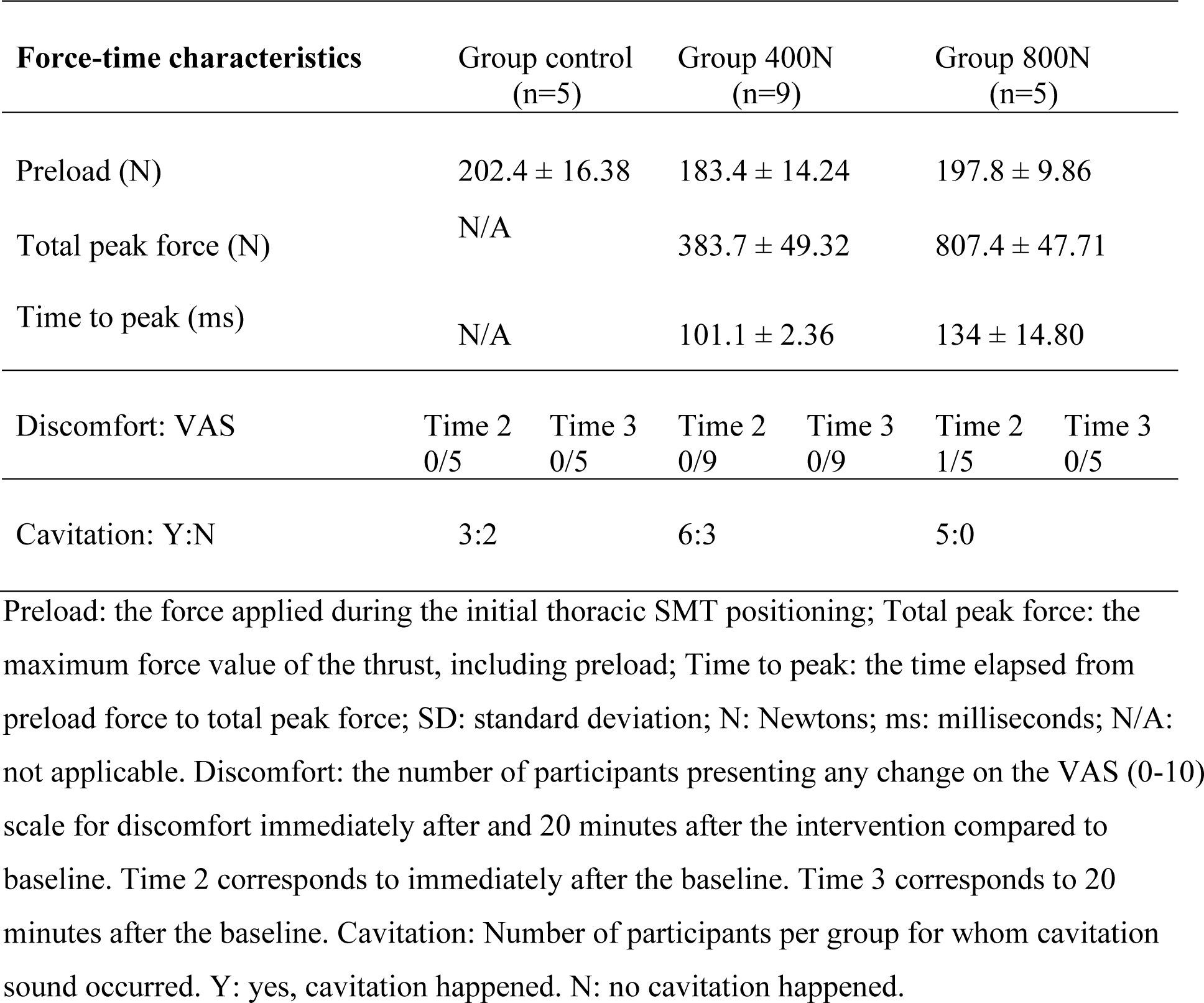
Mean ± SD of the force-time characteristics of the studied groups, number of participants presenting discomfort and cavitation after the correspondent interventions.

### Cavitation and Discomfort

The perception of the participant’s cavitation, characterized by an audible sound during the SMT intervention, was summarized in Table 2. In short, all participants in the target 800N group had thoracic cavitation, while six and three were noted in the target 400N and control group, respectively. None of the participants reported any discomfort after the intervention compared to baseline except one in the target 800N group (Table 2).

### Blood biomarker of oxidative stress and pro-resolution of inflammation

Table 3 presents the mean ± SD values of the pro-oxidants, antioxidants and pro-resolution mediator at baseline, immediately after, and 20 minutes after intervention. Pro-oxidants, antioxidants and resolving D1 parameters were normally distributed: Shapiro-Wilk test (p>0.05). Also, Figures 2 A and B depict a correlation matrix with coefficient values between two variables right after and 20 minutes after intervention, respectively. There was a significant strong correlation between LOOH and TOS (r=0.72; p=0.001) and a trend between TOS and peak force (r=0.42; p=0.073) right after intervention (Figure 2A). In addition, a significant correlation between LPO and peak force (r=0.60; p=0.007), LOOH and TOS (r=0.55; p=0.015) and a negative correlation between TOS and resolving D1 (r= -0.57; p= 0.011) were observed (Figure 2B). No other correlation was statistically significant (p<0.05).

**Figure 2:**
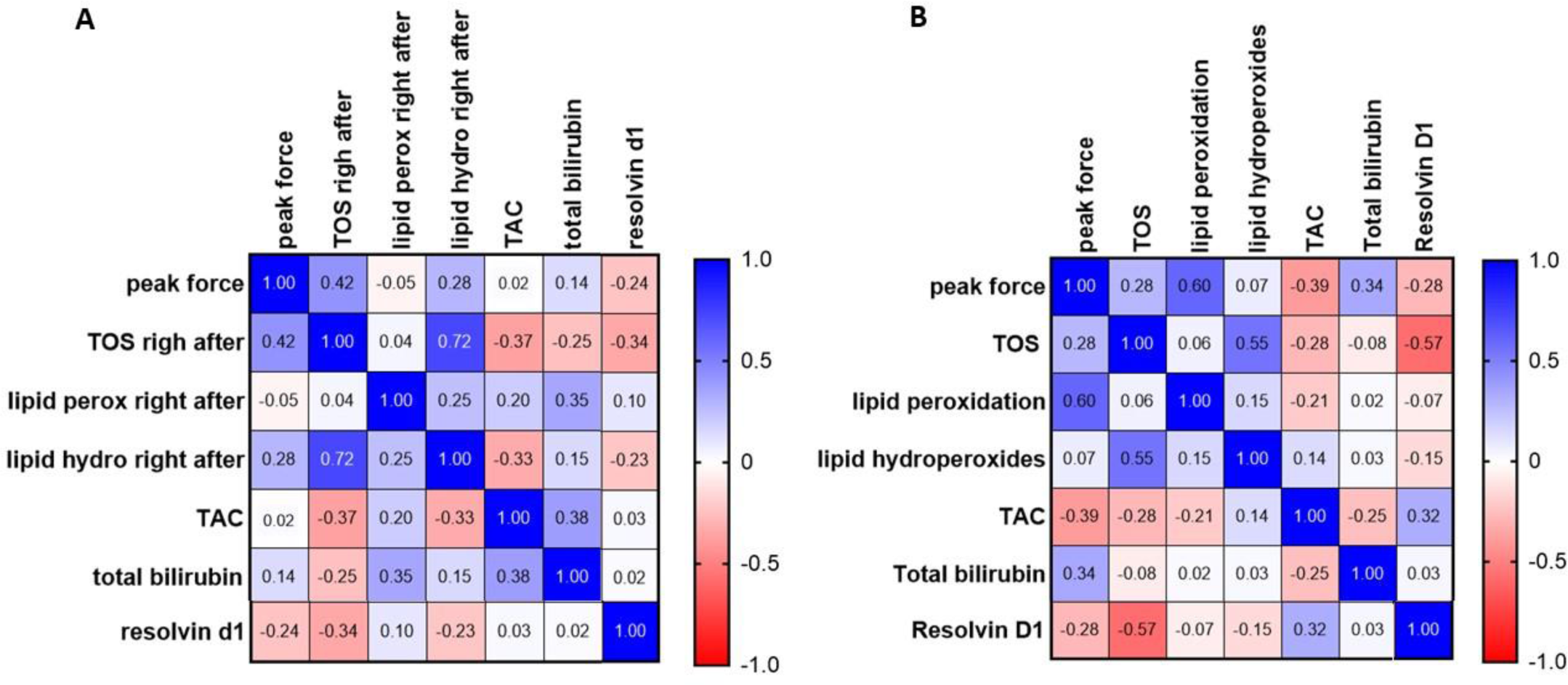
Paired Pearson correlation matrix between peak force and blood biomarkers of oxidative stress and pro-resolution mediator right after intervention (A) and 20 minutes after intervention (B). N= 19. Peak force: maximum force value registered during preload (control group), or maximum force value registered during preload + thrust (target 400N and 800N); TOS: total oxidant status; Lipid perox: (LPO-Lipid peroxidation); Lipid hydro: (LOOH-Lipid Hydroperoxide content); TAC: total antioxidant capacity; Total Bilirubin and Resolvin D1.

**Table 3:**
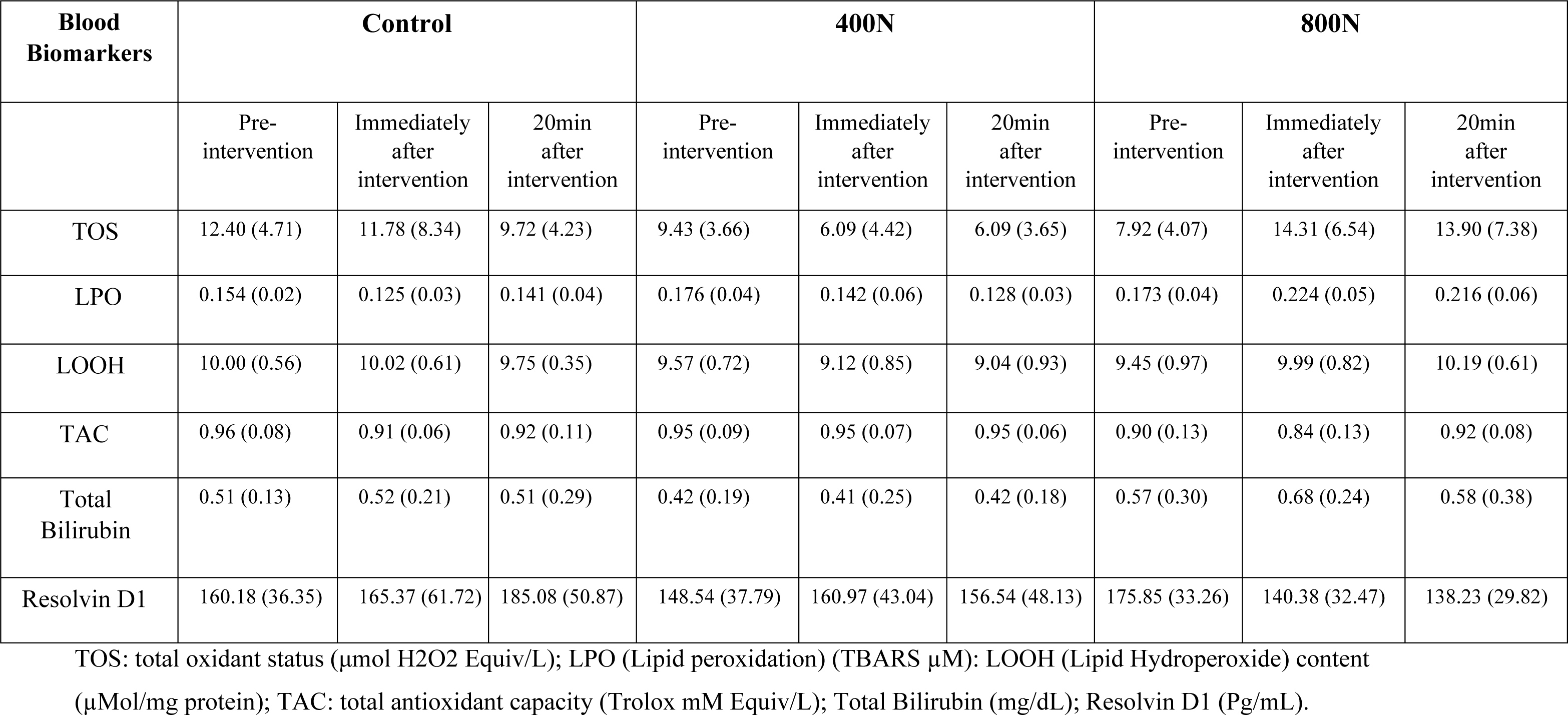
Short-term mean ± (SD) group values of pro-oxidants, antioxidants and resolving D1 in blood in the studied groups.

| Blood Biomarkers | Control |  |  | 400N |  |  | 800N |  |  |
| --- | --- | --- | --- | --- | --- | --- | --- | --- | --- |
|  | Pre-intervention | Immediately after intervention | 20min after intervention | Pre-intervention | Immediately after intervention | 20min after intervention | Pre-intervention | Immediately after intervention | 20min after intervention |
| TOS | 12.40 (4.71) | 11.78 (8.34) | 9.72 (4.23) | 9.43 (3.66) | 6.09 (4.42) | 6.09 (3.65) | 7.92 (4.07) | 14.31 (6.54) | 13.90 (7.38) |
| LPO | 0.154 (0.02) | 0.125 (0.03) | 0.141 (0.04) | 0.176 (0.04) | 0.142 (0.06) | 0.128 (0.03) | 0.173 (0.04) | 0.224 (0.05) | 0.216 (0.06) |
| LOOH | 10.00 (0.56) | 10.02 (0.61) | 9.75 (0.35) | 9.57 (0.72) | 9.12 (0.85) | 9.04 (0.93) | 9.45 (0.97) | 9.99 (0.82) | 10.19 (0.61) |
| TAC | 0.96 (0.08) | 0.91 (0.06) | 0.92 (0.11) | 0.95 (0.09) | 0.95 (0.07) | 0.95 (0.06) | 0.90 (0.13) | 0.84 (0.13) | 0.92 (0.08) |
| Total Bilirubin | 0.51 (0.13) | 0.52 (0.21) | 0.51 (0.29) | 0.42 (0.19) | 0.41 (0.25) | 0.42 (0.18) | 0.57 (0.30) | 0.68 (0.24) | 0.58 (0.38) |
| Resolvin D1 | 160.18 (36.35) | 165.37 (61.72) | 185.08 (50.87) | 148.54 (37.79) | 160.97 (43.04) | 156.54 (48.13) | 175.85 (33.26) | 140.38 (32.47) | 138.23 (29.82) |

TOS: total oxidant status ( $\mu\text{mol H}_2\text{O}_2$ Equiv/L); LPO (Lipid peroxidation) (TBARS $\mu\text{M}$ ); LOOH (Lipid Hydroperoxide) content ( $\mu\text{Mol/mg protein}$ ); TAC: total antioxidant capacity (Trolox mM Equiv/L); Total Bilirubin (mg/dL); Resolvin D1 (Pg/mL).

### Pro-oxidants: TOS

In the present study, we observed an overall decrease in plasma TOS in both control and target 400N groups. In contrast, an 80% and 75% increase in TOS was observed immediately after and 20 minutes after SMT, respectively, compared to baseline. There was an interaction effect between intervention and time [F (4, 32) = 4.04, p=0.009] on plasma TOS. Tukey’s multiple comparison test showed a significant difference between 400N and 800N (MD= -4.75; 95% CI [-9.364 to -0.142], p=0.042) (Figure 3A). In addition, multiple comparison tests showed that compared to baseline, a significant within-group decrease in TOS levels was observed in the target 400N group immediately after (MD=3.63; 95%CI [1.011 to 6.255]; p=0.019) and 20 minutes after intervention (MD=3.38; 95% CI [2.059 to 4.701]; p<0.001 (Figure 3A). Despite a slight decrease in TOS compared to the baseline in the control group or the increase in the target 800N, no further main effect was observed, nor within and between-group difference through pairwise post-hoc test (p>0.05).

**Figure 3:**
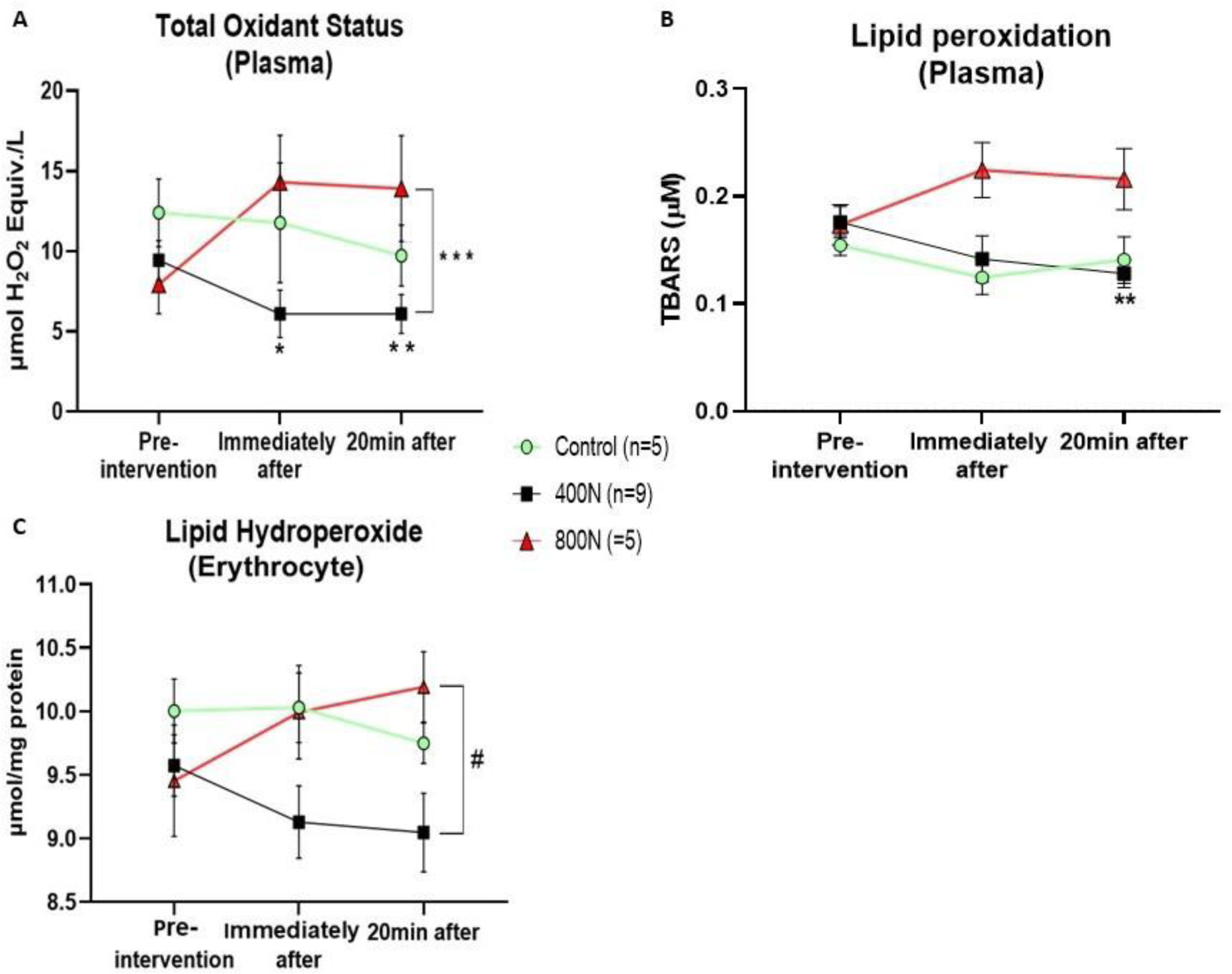
Oxidative stress parameters of plasmatic total oxidant status (A), lipid peroxidation (B), and lipid hydroperoxide (C) in erythrocytes. Measurements were assessed pre-intervention (baseline), immediately after, and 20 minutes after intervention. Control intervention (preload only); 400N (target 400N thoracic SMT); 800N (target 800N thoracic SMT). SMT: posterior to anterior thoracic spinal manipulation. Values are presented as mean and standard error. Significance was set as an alpha of 0.05. * Denotes significance between baseline and immediately after the intervention. ** Denotes significance between baseline and 20 minutes after intervention. *** denotes significance between 400N and 800N groups. # denotes significance between 400N and 800N groups 20 minutes after intervention.

### Pro-oxidants: LPO

Plasma LPO measured by TBARS had about a 20 and 30% decrease over time in the control and target 400N, respectively, compared to the baseline. In contrast, an increase of approximately 30% over time was observed in the target 800N compared to the baseline. Analysis of plasma LPO by mixed-effect model yielded a significant main effect of treatment [F (2, 16) = 4.66, p=0.025] and interaction effect (intervention x time) [F (4, 32) = 3.10, p=0.028]. Tukey’s multiple comparison tests showed that the decrease of LPO 20 minutes after 400N SMT was statistically significant compared to baseline level (MD=0.046; 95% CI [0.014 to 0.078]; p=0.007) (Figure 3B). No other main effect was observed, nor within and between-group statistical difference (p>0.05).

### Pro-oxidants: LOOH content

An overall increase of 15% in erythrocyte LOOH content was observed in the target 800N, while a decrease of approximately 15% was observed in the target 400N group. Mixed-effect model analysis revealed an interaction effect between intervention and time in the LOOH content in erythrocytes [F (4, 32) = 2.86, p=0.038]. Tukey’s multiple comparison tests indicated a statistical difference between the 400 and 800N groups 20 min after intervention (MD= -1.147; 95% CI [-2.263 to -0.030]; p=0.044) (Figure 3C). No other main effect was observed, nor within and between-group statistical difference (p>0.05).

### Antioxidants: TAC

The Mixed-effect model test observed no main effect of the intervention [F (2, 16) = 1.11, p = 0.325], time [F (2, 16) = 1.02, p = 0.358] or interaction effect between intervention and time [F (4, 32) = 0.57, p = 0.685] (Figure 4A). Despite the common sense that the multiple comparison tests are best done following the omnibus test’s rejection of the null hypothesis, the outputs from the two tests may only sometimes be concordant since they assess different aspects of the data. While the multiple comparisons test aims to determine if there is a mean difference using a pairwise comparison, the omnibus test compares the variation within each group to the variation of the mean of each group to test if the null hypothesis can be rejected.^32^ Thus, when a Tukey’s multiple comparison test was conducted, there was evidence of a decrease of TAC immediately after 800N compared to its baseline (MD=0.060; 95% CI [0.008 to 0.112]; p = 0.030) (Figure 4A). No other within nor between-group statistical difference was observed (p>0.05)

**Figure 4:**
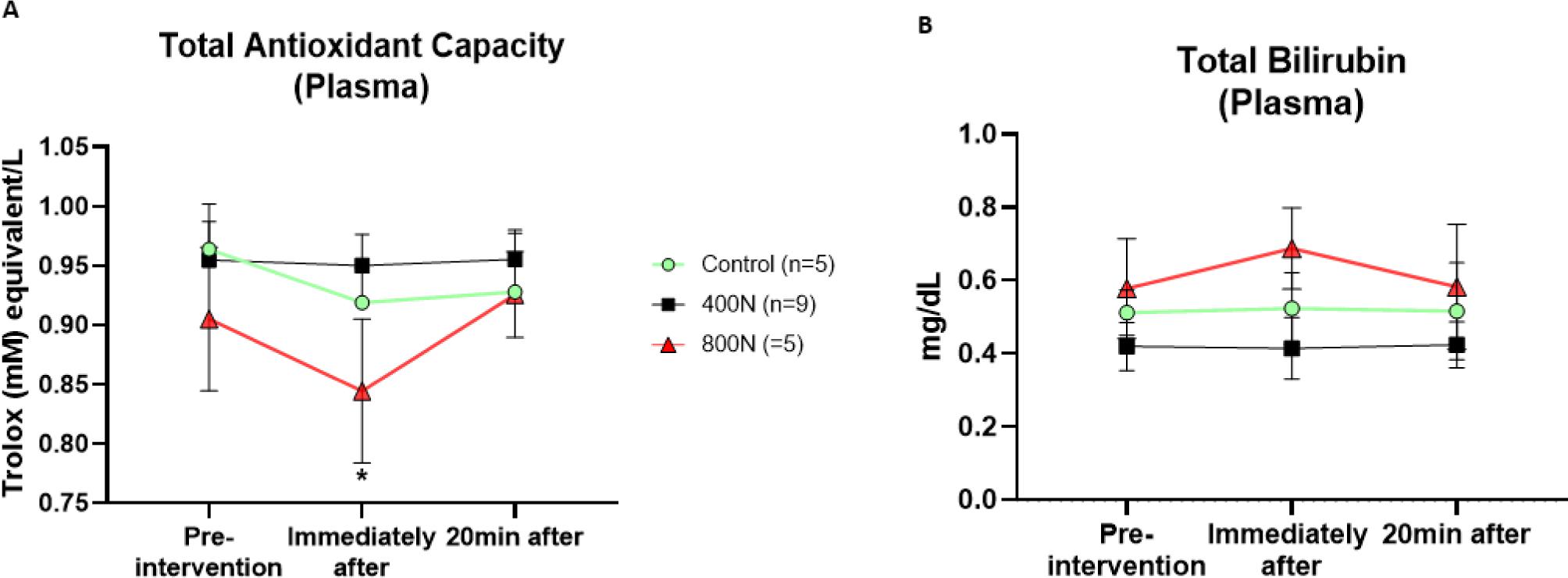
Plasmatic antioxidant parameters of total antioxidant capacity (A) and total bilirubin (B). Measurements were assessed pre-intervention (baseline), immediately after, and 20 minutes after intervention. Control intervention (preload only); 400N (target 400N thoracic SMT); 800N (target 800N thoracic SMT). SMT: posterior to anterior thoracic spinal manipulation. Values are presented as mean and standard error. Significance was set as an alpha of 0.05. * Denotes significance between baseline and immediately after the intervention.

### Antioxidants: Total Bilirubin

No main effect of the intervention [F (2, 16) = 0.96, p = 0.401], time [F (2, 16) = 0.06, p = 0.926] nor interaction effect between intervention and time [F (4, 32) = 0.46, p = 0.759] was observed in the assessment of total bilirubin (Figure 4B). No other within nor between-group statistical difference was observed (p>0.05).

### Pro-Resolution Mediator: Resolvin D1

There was an interaction effect between intervention and time [F (4, 32) = 2.99, p=0.032] on plasma Resolvin D1. Multiple comparison tests showed that Resolvin D1 levels significantly decreased 20 minutes after 800N SMT compared to baseline (MD=37.62; 95% CI [9.462 to 65.80]; p=0.019) (Figure 5). No other main effect was observed, nor within and between-group statistical difference (p>0.05).

**Figure 5:**
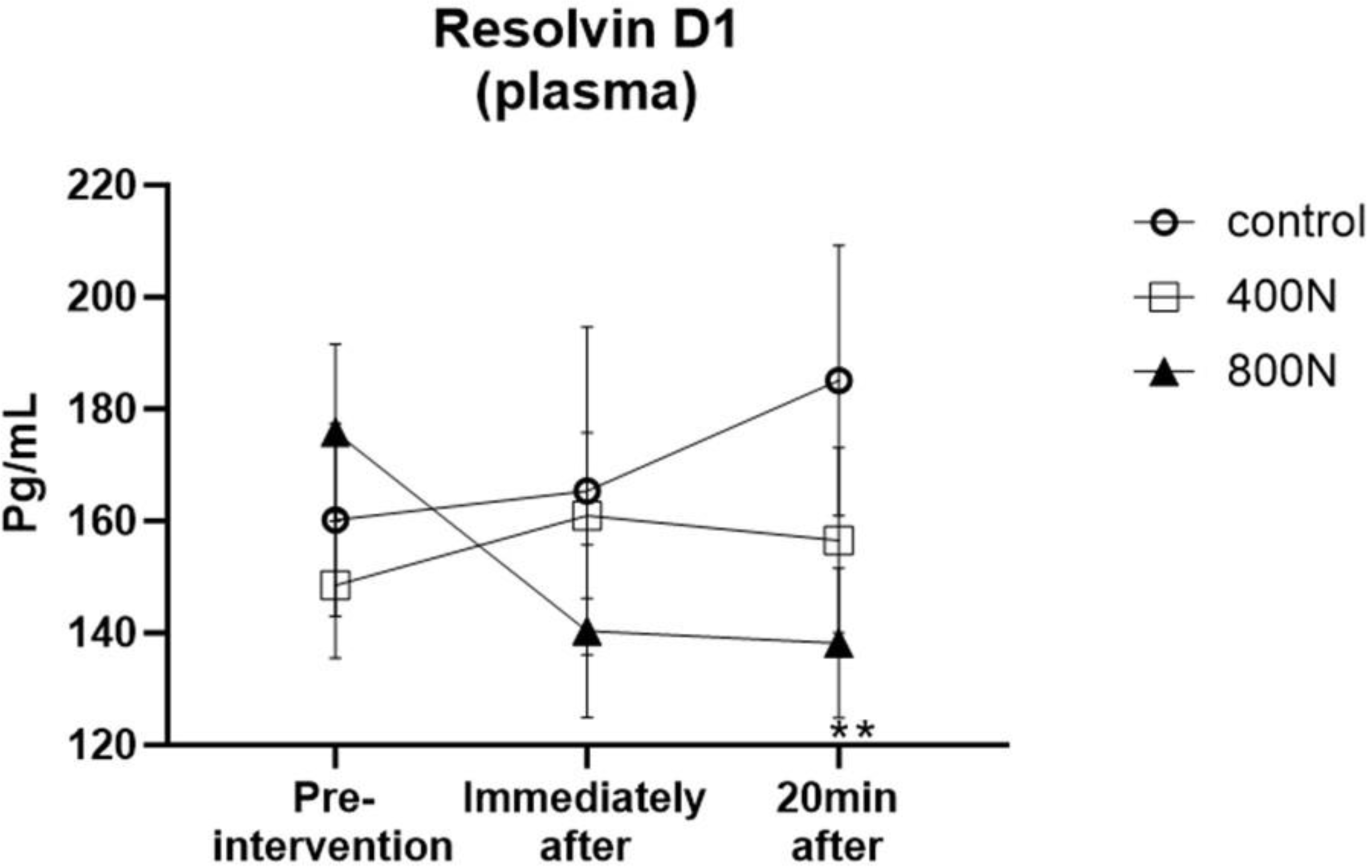
Lipid-derived resolution mediator resolving D1 level in plasma. Measurements were assessed pre-intervention (baseline), immediately after, and 20 minutes after intervention. Control intervention (preload only); 400N (target 400N thoracic SMT); 800N (target 800N thoracic SMT). SMT: posterior to anterior thoracic spinal manipulation. Values are presented as mean and standard error. Significance was set as an alpha of 0.05. ** Denotes significance between baseline and 20 minutes after intervention.

## Discussion

The present study examined the short-term effect of a single thoracic SMT with different force-time characteristics (force magnitudes) on systemic blood oxidative stress biomarkers and lipid-derived pro-resolution mediators in healthy subjects. The essential observation from this study was the likely contrasting effects between the employed target 400 and 800N SMT on the biomarkers assessed. Target 800N SMT seemed to lead to an increase of plasma and erythrocyte pro-oxidants (TOS, LPO and LOOH) with a parallel decrease in plasma TAC and resolving D1; in contrast, target 400N SMT seemed to decrease plasma and erythrocytes pro-oxidants TOS, LPO and LOOH, respectively with no changes in antioxidants nor resolvin D1 levels. These observations support our hypothesis that different SMT force-time characteristics may trigger contrasting effects on the assessed systemic pro-oxidant, antioxidant biomarkers, and pro-resolution mediators (resolving D1). In addition, it builds on our previous findings of a panel of inflammatory cytokines presenting opposing behaviour upon target 400 and 800N SMT thoracic SMT.^4^

The oxidant/antioxidant system is a complex and fined-tune system of *in vivo* physiology. Systemic redox signalling biomarkers have promising clinical utility in clustering patients according to biomarker characteristics. Previous studies on patients with cardiovascular disorders have shown that systemic redox signalling biomarkers were associated with the risk of developing cardiovascular events in post-menopause females and with poorer whole-body aerobic capacity, with direct implications in physical exercise tolerance in patients with chronic heart failure.^33,34^ In chronic MSK conditions such as osteoarthritis and low back pain, studies have shown that inflammatory cytokines and redox signalling biomarkers, which play a crucial role in regulating inflammatory responses, may be used to identify patients with a higher likelihood of presenting lower physical function, higher pain intensity, greater pain catastrophizing, and multiple painful areas (widespread pain).^16,35^ Thereby, treatment strategies that aim to modify these biomarkers are of great relevance. In this regard, the findings from the present study are of great potential given the 400N and 800N SMT’s ability to modify systemic oxidative stress biomarkers, which is in agreement with pre-clinical and clinical evidence that SMT can modulate oxidants and antioxidant biomarkers while improving pain and physical function.^7,9,18^ Despite not being in a clinical population, our present study reinforces previous findings on the modulatory role of SMT on systemic oxidative stress biomarkers.

While several parameters can be used to estimate redox signalling biomarkers, we assessed 1) TOS to estimate the total oxidant content in the plasma sample, 2) LPO by measuring its final product formation after the action of oxidants on cellular membrane polyunsaturated fatty acids, and 3) LOOH in erythrocytes, which is an intermediate product of lipid oxidation.^36,37^ Our findings on oxidants showed that single thoracic SMT of 400N seemed to provide a short-term reduction in TOS compared to baseline and the target 800N group (Figure 3). Similarly, a decrease in the LPO and LOOH levels was observed in the 400N group. Such a decrease likely allowed the within-group difference between baseline and 20 minutes in the LPO levels and between 400N and 800N in the LOOH.

In the opposite direction to what was observed in the 400N group, the 800N SMT induced a slight increase in all systemic oxidant biomarkers compared to baseline. These results suggest that only 800N increases oxidative stress in plasma and erythrocytes after SMT. Interestingly, peripheral blood mononuclear cells from healthy humans presented an initial reduction in mitochondrial activity after an oxidative challenge. Still, the mononuclear cells restored the physiological activity levels at three hours.^38^ Since we did not evaluate oxidative biomarkers longer than 20 minutes, we cannot exclude the possibility that mononuclear cells are involved in the changes in oxidative stress biomarkers after 400N and 800N SMT.

Our preliminary findings suggest that the target 800N SMT seemed to rise in systemic oxidant biomarkers in the short term. Increased oxidants after 800N SMT may be considered harmful and related to adverse events. For instance, after acute intense exercise in untrained healthy participants, an increase in systemic oxidants such as LPO is associated with the perception of exercise-induced pain in the short-term post-exercise recovery period.^39^ In addition, in patients with chronic fatigue syndrome, an accentuated LPO was observed with a parallel decrease in non-enzymatic antioxidants in both a resting state and an acute bout of exercise.^40–42^ In the present study, only one participant in the target 800N group reported an increase in the VAS scale from zero to one immediately after the intervention. Also, despite the increase of oxidants, the values observed did not correspond to systemic levels observed in pathological disorders such as cancer and multiple sclerosis or fibromyalgia and osteoarthritis.^43–46^ Therefore, the heightened oxidant levels observed in the short-term after target 800N SMT might not be the main drivers underlying the perception of the discomfort or adverse events up to 20 minutes after SMT, nor be deleterious/pathological. In addition, a pre-post study using massage therapy as an intervention reported increased levels of oxidants and decreased antioxidants in healthy young participants.^47^ Such a pattern of oxidants/anti-oxidants has also been described after a bout of physical activity.^37,39^ The transient but not persistent, enhanced redox signalling biomarkers after acute physical activity are crucial in driving muscular, neuroimmune and other systemic adaptations, paving the foundations of regular exercise benefits in the neuro-immune system and pain.^10^ Therefore, we speculate that target 800N may lead to a short-term, not persistent, transient increase of oxidative stress, which may have an adaptive functional role as a secondary messenger and in activating transcription factors to enhance cellular survival instead, similar to what has been shown in other therapeutic strategies, such as regular physical activity.^48^ However, further research is needed to determine whether the increase of oxidative stress is a transitory response followed by 800N SMT and to understand this complex relationship between oxidative stress parameters and mechanical loadings of mechanotherapies.

Parallel to changes in oxidant biomarkers, 800N total peak force SMT significantly decreased plasma TAC. Human plasma is endowed with an array of antioxidant defence mechanisms. TAC assay is widely used to estimate the global enzymatic (e.g., superoxide dismutase, catalase and selenium-dependent glutathione peroxidase) and non-enzymatic (e.g., ascorbate, urate, a-tocopherol, bilirubin, albumin), antioxidant components of a sample.^36,49,50^ Since 800 SMT triggered a decrease in TAC levels, the antioxidant system was possibly required to counteract the increase of oxidants observed after 800N. Organisms maintain oxidative homeostasis through a sophisticated regulatory system that includes enzymatic and non-enzymatic antioxidants.^51^ Since the technique used in the present study to assess TAC measures mainly albumin and uric acid, but ascorbic acid, α-tocopherol and bilirubin are also measured, we would expect a change in bilirubin levels after 800N SMT.^52^ However, no statistical difference after 800N SMT was observed on bilirubin levels despite the slight elevation immediately after the intervention. Total bilirubin functions as a plasma non-enzymatic antioxidant, inhibiting nicotinamide adenine dinucleotide phosphate (NADPH) oxidase activity - a significant source of oxidants and free radicals - and synergistically interacting with other antioxidants to inhibit lipid oxidation.^53,54^ Thus, future studies by our group will better assess the relationship between force magnitude and bilirubin and other antioxidant levels after 800N SMT.

For the first time, the effects of SMT on lipid-derived pro-resolvin molecules were demonstrated. We observed a statistical decrease in resolving D1 in the target 800N group compared to its baseline, while no changes were observed in the target 400N or the control group. Resolvin D1, a novel specialized pro-resolving lipid mediator, contributes to the resolution of inflammation by reducing neutrophil trafficking to the injured site, diminishing oxidative stress and promoting the clearance of tissue debris and apoptotic cells.^20^ Evidence also supports the role of resolvin D1 in controlling the exacerbation of afferent nerve activation, such as nociceptors, during peripheral sensitization and mechanical hyperalgesia through action on different TRP afferent receptors.^55^ Thus, the decrease in resolving D1 after 800N SMT may be related to the parallel increase in oxidants such as TOS, negatively correlated with resolving D1. However, further basic science and clinical studies are warranted to understand further how SMT’s force-time characteristics relate to pro-resolution mediators and the utility of resolvins in spine pain and other cMSK conditions.

How could SMT alter oxidative stress and resolvin D1 levels? Even though it is early to speculate, it has been suggested that mechanical stimuli have the potential to be translated into mechanochemical signals through mechanotransduction.^56^ At a cellular level, such signals can modulate local cellular factors in their respective plasma membrane or propagate the signals from the cellular plasma membrane to the nucleus, altering intracellular signalling biochemistry and gene activity.^56^ Several endogenous sources can generate redox signalling biomarkers, such as fibroblasts, neutrophils, monocytes and macrophages, vascular endothelial cells, neurons, and glial cells. ^57–59^ However, muscle cells, neutrophils, monocytes, and macrophages are significant candidates contributing to the systemic levels.^33^ In healthy individuals, the responses of both polymorphonuclear neutrophils and monocytes were significantly higher after SMT (compared to baseline) and significantly higher than in sham or soft-tissue treated subjects.^60^ In activated immune cells, metabolic reprogramming during immune responses directly produces excessive cytosolic and mitochondrial reactive oxygen species.^61^ Resolvin D1 rescued macrophages from oxidative stress-induced apoptosis during efferocytosis, promoting resolution of inflammation.^62^ Thus, immune cells such as monocytes and macrophages may be the link between the changes induced by SMT on oxidative stress biomarkers and resolving D1 levels. However, this putative relationship should be further explored.^73^

Although our results suggest that SMT force magnitude influences blood biomarkers of oxidative stress, this study has some limitations. The first limitation is the small sample size. Due to the COVID-19 global pandemic, participant recruitment was limited, and our pre-established target sample size could not be reached. This weakened the analyses of variances within and between groups. Despite this limitation, the results of variance analysis from this preliminary study may help to guide future studies about SMT force-time characteristics and oxidative/antioxidant parameters. The second limitation was that only a few oxidative/antioxidant parameters were assessed. A recent study highlighted that a single redox signalling marker is of limited diagnostic and prognostic value and is insufficient to determine the extent and the implication of the oxidative stress of a given tissue/organ^36^. Therefore, a more comprehensive range of assays is recommended. The third limitation is that although no perception of discomfort/adverse event was reported, participants were only followed up to 20 minutes after SMT. It is known that most adverse events occur within the first 48 hours after SMT^63^. Therefore, we cannot discount the possibility that adverse events occurred after the data collection period. Also, most participants had previous experience with chiropractic care and, thus, were not naïve to SMT. Being familiar with the treatment may mitigate perceived discomfort after intervention. Fourth, we conducted the study with young and healthy participants, limiting the findings’ interpretability to the clinical population. However, exploring the force-time characteristics in non-clinical participants allows us to determine which characteristics and outcomes are modifiable by different SMT force-time characteristics while providing highly valuable research direction and baseline values to inform future research in clinical populations. Fifth, we confined our determinations to oxidant/antioxidant biomarkers in systemic blood. Future studies are needed to determine a more comprehensive understanding of the SMT force-time characteristics and redox signalling biomarkers in several other cell types (e.g., muscle, white blood cells). Despite the limitations, this study explored a novel research question to explore how distinct SMT mechanical loadings influence systemic biomarkers of oxidative stress using a reliable, valid and reproducible methodology (FSTT®) to quantify the force-time characteristics during manual therapy interventions (i.e., SMT).^64,65^ Unveiling the physiological events associated with mechanotherapies in a healthy cohort is a first step before studying the clinical population.

## Conclusion

The findings from the present study showed that while 400N seemed to decrease systemic TOS and lipid peroxidation biomarkers, 800N appeared to increase these parameters with further reduction in plasma TAC and resolving D1. Thus, our findings may indicate that different SMT force magnitudes applied to the thoracic spine may have a contrasting effect on the systemic redox signalling biomarkers and pro-resolution mediators in healthy participants. Future studies are required to determine whether similar findings are seen in individuals with cMSK disorders such as CPLBP and elucidate the clinical value of SMT’s force-time characteristics on oxidative stress and pro-resolution mediators.

## Acknowledgement

We thank Dr. Stephen Injeyan for sharing his expertise and thoughtful insights during study design and data collection and Isabella Magnani and Steven Tran for their technical support during data collection and analysis.

